# Spatial Correlation as an Early Warning Signal of Regime Shifts in a Multiplex Disease-Behaviour Network

**DOI:** 10.1101/195388

**Authors:** Peter C. Jentsch, Madhur Anand, Chris T. Bauch

## Abstract

Early warning signals of sudden regime shifts are a widely studied phenomenon for their ability to quantify a system’s proximity to a tipping point to a new and contrasting dynamical regime. However, this effect has been little studied in the context of the complex interactions between disease dynamics and vaccinating behaviour. Our objective was to determine whether critical slowing down (CSD) occurs in a multiplex network that captures opinion propagation on one network layer and disease spread on a second network layer. We parameterized a network simulation model to represent a hypothetical self-limiting, acute, vaccine-preventable infection with shortlived natural immunity. We tested five different network types: random, lattice, small-world, scale-free, and an empirically derived network. For the first four network types, the model exhibits a regime shift as perceived vaccine risk moves beyond a tipping point from full vaccine acceptance and disease elimination to full vaccine refusal and disease endemicity. This regime shift is preceded by an increase in the spatial correlation in non-vaccinator opinions beginning well before the bifurcation point, indicating CSD. The early warning signals occur across a wide range of parameter values. However, the more gradual transition exhibited in the empirically-derived network underscores the need for further research before it can be determined whether trends in spatial correlation in real-world social networks represent critical slowing down. The potential upside of having this monitoring ability suggests that this is a worthwhile area for further research.

## 1. Introduction

Vaccine-preventable infectious diseases continue to impose significant burdens on populations around the world [1]. Access to vaccines remains a significant barrier to providing more widespread protection against infectious diseases. However, a growing obstacle to infection control is vaccine refusal, which can have a large effect on disease prevalence. For instance, the drop in vaccine coverage after Andrew Wakefield’s fraudulent 1998 paper about the mumps-measles-rubella vaccine reduced MMR coverage to as low as 61 % in some areas of the United Kingdom [2].Lower vaccine coverage caused larger measles outbreaks in the years following the publication of the Wakefield paper [3][4]. Elimination of polio in Africa was similarly interrupted when a rumor that the vaccine could cause infertility or HIV infection began spreading in 2003, when leaders of three states in north-central Nigeria boycotted the vaccine until it could be tested independently. The impasse was not resolved until the following year, a time period during which these states accounted for over 50% of polio cases worldwide [5, 6]. Vaccine refusal and hesitancy are also common for influenza vaccine, with non-vaccinators citing concern for side effects, lack of perception of infection risk, and doubts about vaccine efficacy as reasons to not become vaccinated [7].

Simple differential equation models such as the Kermack-McKendrick SIR (susceptible-infected-recovered) model published in 1927 (originally formulated as an integro-differential equation) [8], allow us to characterize useful measures such as the expected number of new infections caused by each infection, and are readily fitted to epidemiological data. Classical infection transmission models such as the Kermack-McKendrick model assume that members of the population mix homogeneously. However, in many situations, infection transmission through a network–where individuals are nodes and contacts through which infection may pass are edges–are a more accurate description of infection dynamics [9]. Networks tend to be analytically intractable and therefore agent-based models are often used to simulate networks. Agent-based simulations on networks allow us to specify complex individual node behavior in a natural way. One of the most ambitious examples of these is the Global-Scale Agent Model, which models the daily behavior and relationships of 6.5 billion people using worldwide GIS data[10]. However, agent-based network simulations have also been studied in the context of nonlinear interactions between disease dynamics and individual behaviour concerning vaccines and contact avoidance [11, 12, 13, 14, 15].

The trajectory that an infection takes as it moves through a population is heavily influenced by the spread of health information between individuals, so more sophisticated models of disease spread often combine disease dynamics and social dynamics. The coupled interactions between individual behaviour and disease dynamics have been modelled under various frameworks and placed under various rubrics including: epidemic games [16], coupled behaviour-disease models [12, 17, 18], socio-epidemiology, economic epidemiology and behavioural modeling [19]. A more recent trend in epidemiological modeling is to abstract these two subsystems into (1) an information transfer network through which information flows between individuals, and (2) a separate physical disease transmission network. A system where each node is part of two or more different networks is called a multiplex network, and is a natural way to implement a coupled disease-behaviour system [20, 18]. For instance, the simultaneous spread of disease and disease awareness over adaptive multiplex networks with scale-free degree distributions has been studied [21]. Similarly, a three layer network to model the diffusion of infection, awareness, and preventative measures along different contact networks was found to reasonably approximate empirical influenza data[22]. Similar approaches consider coupled human and ecological dynamics, which present the opposite problem of species that humans wish to preserve instead of eradicate [23, 24, 25, 26].

The nonlinear coupling between disease and social processes creates feedback loops between infection prevention mechanisms and disease spread. Nonlinear feedback in other complex systems such as from solid state physics and theoretical socio-ecology has often been shown to yield critical transitions [27, 28, 26]. A critical transition is defined as an abrupt shift from an existing dynamical regime to a strongly contrasting (and sometimes unfavourable) dynamical regime as some external parameter is pushed past a bifurcation point [29, 30]. Fortunately, critical transitions (and other regime shifts associated with a bifurcation where the dominant eigenvalue of the Jacobian matrix around the equilibrium approaches zero) often exhibit characteristic early warning signals beforehand that allow these shifts to be predicted [31, 32, 30]. Critical slowing-down (CSD) based indicators were one of the first early warning signals to be studied. CSD occurs because the speed with which a system responds to perturbations slows as it approaches bifurcations where the magnitude of the dominant eigenvalue of the Jaacobian approaches zero at the bifurcation point. Since nearly all systems in the real world are subject to perturbations, the lag-1 autocorrelation of a time series can be used as a relatively universal (or at least potentially common) indicator of CSD. Lag-1 autocorrelation appears to be a robust statistic and has been shown to be present in predicting catastrophic bifurcations in complex real world systems such as the global climate[33], human nervous systems[34], and stock markets[35].

The discrete fourier transform (DFT) of a network is another example of a CSD-based early warning signal. Under some assumptions, the WeinerKinchin Theorem shows that we can use the discrete Fourier transform (DFT) to measure spatial correlation in system state, and this has been shown to work in some ecological applications [36] [37]. Lag-1 spatial correlation can in some cases provide a better early warning signal than time-domain methods, because”a spatial pattern contains much more information than does a single point in a time series, in principle allowing shorter lead times” before the critical transition occurs [38, 31]. This observation has been corroborated in three ecological dynamical systems[31].

Early warning signals of regime shifts in coupled behaviour-disease networks have received relatively little attention in the literature on modelling interactions between disease dynamics and human behaviour. This appears to be a significant knowledge gap because early warning signals for vaccine scares could help public health anticipate widespread vaccine refusal and prepare for outbreak response in advance, as well as build efforts to improve trust between the public and the health authorities. In this paper we use an agent-based model on a two-layer multiplex network to simulate the coupled disease dynamics of a vaccine-preventable infection and social dynamics of vaccination in a population. We show that spatial correlation can be used as an early warning signal for regime shifts in this system on most (but not all) network topologies. In the next section we discuss the model structure and methods of analysis, followed by a section on results and finally a discussion section.

## 2. Methods

### 2.1 Simulation

Our agent-based model simulated a population of 10,000 individuals (nodes), where every node belongs to two different connectivity networks: a transmission network and a social network. In the transmission network, each node is connected to other nodes from which they can contract infection. Two nodes are linked in the social network if they can be influenced by one another’s opinions on vaccination. These networks were simulated as fixed graphs upon which stochastic processes occurred, with a variety of degree distributions and average path lengths.

We modelled a hypothetical acute, self-limiting infection with rapidly waning natural immunity Each node on the physical layer is in one of four possible states: susceptible (*S*), infected (*I*), recovered (*R*), or vaccinated (*V*). Each node on the social layer also has an opinion on the vaccine: they are either a non-vaccinator (*η*), or a vaccinator(*v*). We will denote the the biological state of a node *v* by *B*(*v*), and the opinion of a node *v* by Θ (*v*). The transmission network is a graph denoted by *T* (*V, E*_*T*_), and the social network is a graph denoted by *O*(*V, E*_*O*_). We assume that they share the same set of vertices *V* although this assumption could be relaxed in future work. The set of nodes in the neighbourhood of *v* is *adj*_*T*_ (*v*) or *adj*_*O*_(*v*) for the transmission and the social network respectively.

The algorithm used to simulate the social and transmission processes used discrete timesteps. At each time step, for each *v ∈ V*:

- If *B*(*v*) = *I*, then for all *u ∈adj*_*T*_ (*v*) such that *B*(*u*) = *S* and θ(*u*) ≠ *v*, set *B*(*u*) = *I* with probablility *p* (infection event)
- If *B*(*v*) = *I*, let *B*(*v*) = *R* with probability *r* (natural recovery event)
- If *B*(*v*) = *R*, set *B*(*v*) = *S* with probability γ (loss of immunity event)
- If *B*(*v*) = *S*, set *B*(*v*) = *I* with probability *σ* ≪ 1 (case importation event)
- Choose some node *u ∈ adj*_*O*_(*v*) uniformly at random. If Θ (*v*) */*≠ Θ (*u*), then *P* (*η→ν*) = ϕ (*E*_*V*_ *E*_*N*_), and *P* (*ν→η*) = 1 ϕ (*E*_*V*_ *E*_*N*_)

where

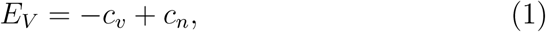

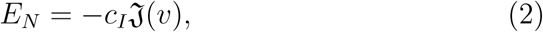

where ϕ is a sigmoid function such that ϕ (*∞*) = 1, ϕ (*∞*) = 0, ϕ (0) = 0.5 as described in previous models (opinion change event) [39]. In our implementation, 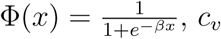 is the perceived cost of vaccination (due to infection risks), *c*_*I*_ is the perceived cost of infection (due to infection risks), *β* controls the steepness of the sigmoid function, and j(*v*) = *|{u ∈ adj*_*T*_ (*v*): *B*(*u*) = *I}|* is the number of infected nodes adjacent to *v* in the transmission network. *c*_*n*_ represents some outside incentive that a person might have for vaccinating, such as peer approval, school admission requirements, or tax incentives. Normalizing both payoff equations by *c*_*I*_ yields

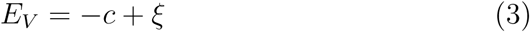

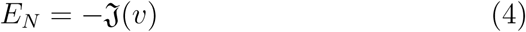

where *c* is the ratio of perceived vaccine risk to perceived disease risk,*c* and 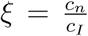 is the ratio of the vaccination incentive to the perceived disease risk. Since changes in perceived vaccine risk are controlled through changes in *c*, we will vary *c* in our analysis. We assume the vaccine is perfectly efficacious.

- With probability *E, v* changes opinions (random opinion change event). That is, if Θ (*v*) = *v*, set Θ (*v*) = *η* and vice-versa.
- If the opinion of a node changes to vaccinator, then their physical state changes to immunized immediately. If they change back to a nonvaccinator, they become susceptible immediately.

We applied synchronized updating to the network: the change in state resulting from each rule is stored and applied after every rule is checked, so the order of the above steps does not matter.

The result of these rules is a feedback loop where, depending on the relative costs of vaccination and infection, the population tends not to exhibit a mixture of strategies except near the critical values of *c*. When *c < ξ*, the payoff to vaccinate *E*_*V*_ is positive and thus exceeds the payoff not to vaccinate *E*_*N*_ which always obeys *E*_*N*_ ≤0. In this case, in the limit as *β → ∞*, all nodes are therefore vaccinators and the infection dies out. However, when *c > ξ* and thus *E*_*V*_ *<* 0, the disease-free equilibrium destabilizes since *E*_*N*_ *≈* 0 in the absence of sustained transmission. In general, since the vast majority of nodes do not have infected neighbours at the disease-free equilibrium, there is a rapid shift in the population to non-vaccinator opinions as well as epidemic outbreaks. For larger values of *β*, the function controlling the opinion-switching as a function of the payoff difference between vaccinator and non-vaccinator strategies is steeper, and the population transition from non-vaccinator to vaccinator strategies is therefore sharper, yielding a critical transition. However, we will use the more general term ‘regime shift’ throughout this paper, since the transition can be made more or less abrupt by changing the value of *β*.

### 2.2 Early Warning Signal Analysis

As the system approaches a regime shift, the dominant eigenvalue of the underlying dynamical system will approach zero. Therefore, it will take longer for the system to recover from perturbations to the steady state. In a spatially extended population, this will increase population heterogeneity as small clusters of non-vaccinators begin to emerge, as well as causing longrange correlations to develop across the network in a detectable way [31]. This development is reflected by an increase in a statistic called the lag-1 spatial correlation (lag-1 SC). We used Moran’s I to measure the lag-1 SC of non-vaccinators as described in [40]. Moran’s I is widely used to calculate the spatial correlation for early warning signals [41, 42, 43].

Let *G* = (*V, E*) be a graph with *n* nodes, *adj*(*v*) be the set of vertices adjacent to *v*, and *f* (*v*) be a binary function such that *f* (*v*) = 1 if *v* is a vaccinator, and *f* (*v*) = 0 otherwise. We define Moran’s I at lag-1, called *M* to prevent confusion with the Infected state, as:

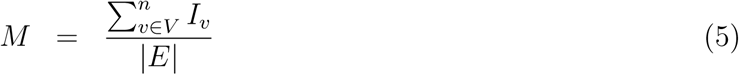

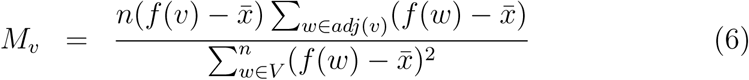

where 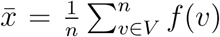 is the fraction of vaccinators in the network. Far from the regime shift, we have that 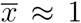 and *f* ≈ 0 for all nodes, thus *I* ≈ 0. However, as resilience to perturbations declines close to the regime shift, the population become more heterogeneous. This causes 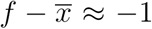 in correlated non-vaccinator clusters, thus *I* increases.

For each realization, the simulation was run long enough for the spatial correlation to stabilize (3500 timesteps), and the equilibrium value was calculated as the average of the next 500 measurements. The equilibrium lag-1 SC was obtained for 100 realizations of the simulation, and these values were averaged to obtain a data point for every value of *c*. The social network and the transmission network are always both the same type of network, but independently generated.

### 2.3 Parameter Values

Baseline parameter values appear in Table 1. The parameter values were chosen to qualitatively represent a hypothetical acute-self limiting infection with waning natural immunity, such as the case of meningococcal infection, influenza or pertussis [44, 45, 46, 47]. The value for γ corresponds to a mean duration of infection of 14 days, the value for γ corresponds to losing natural immunity after two years, and the value for *σ* corresponds to a case importation event in the network once every two months. We conduct univariate sensitivity analysis with respect to *r* and *σ*, since they are important parameters governing the natural history of the infection. For the baseline parameter values, *ξ* is set to zero without loss of generality. The value of *c* will be varied in the analysis of early warning signals. *∈>* 0 is required to prevent the population from fixating on one of the two strategies. To initialize each stochastic realization, one randomly chosen node is infected, and each node is a vaccinator with probability 0.5.

**Table 1:** Parameter definitions and baseline parameter values in probability per timestep (unless otherwise stated). One timestep was interpreted to correspond to one day.

| Parameter | Value | Definition |
| --- | --- | --- |
| $p$ | 0.5 | Probability that an infected node infects a given susceptible neighbour |
| $r$ | 0.07143 | Probability that an infected node recovers |
| $\gamma$ | 0.001369 | Probability that a recovered node becomes susceptible |
| $\epsilon$ | 0.001369 | Probability that a node randomly switches their opinion on vaccination |
| $\sigma$ | 0.016666 | Probability of disease reintroduction |
| $\xi$ | 0 | Parameter governing incentive to become vaccinated |
| $c$ | 0.1 | Ratio of perceived risk of vaccine to perceived risk of disease |
| $\beta$ | 1 | Parameter controlling the steepness of $\Phi$ |

### 2.4 Networks

We ran our model on five different networks: Erdos-Renyi [48], Barabasi-Albert [49], square lattice (or grid), Kleinberg small world[50], and ten subsets of a network constructed by the Network Dynamics and Simulation and Science Laboratory (NDSSL), based on GIS data from the city of Portland [51].

An Erdos-Renyi network is simply given a set of nodes *V* and *v, w ∈, V, v* is connected to *w* with some probability *p*. In our Erdos-Renyi network model, we used a connection probability of 0.001, so each node has degree 10 on average.

The Barabasi-Albert model yields networks with a scale-free (or powerlaw) node degree distribution. Starting with a small initial connected network (*V, E*), new nodes are added to *V* one at a time. Where the probability that the new node is connected to an existing node 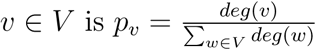. To ensure that the network is always connected, new nodes are also connected to *m* existing vertices, chosen uniformly at random. The Barabasi-Albert networks we used had *m* = 1

Our lattice with *n* = 10, 000 nodes was built as follows: if the nodes are arranged on the integer points of a square 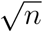 units wide, each node is connected to the nodes within a unit distance up or down (but not both). Because lattice networks are not random, there is no difference between the social and transmission networks and therefore this is effectively not a multiplex network.

The Kleinberg small world network is defined as a square lattice, where additional edges are added between some nodes *v* and *w* with a probability proportional to 1*/d*(*v, w*). The result of this process is a network with a very short average path length. In our implementation, nodes only gain extra edges with 0.1 probability.

The empirically-derived networks from the NDSSL dataset are designed to have some of the properties of a real contact network, being derived from the population of Portland, Oregon. We used a set of ten subnetworks sampled from the NDSSL dataset and constructed in such a way to share the same properties as the original dataset (see Ref. [39] and supplementary appendix for details). The subnetworks had an average path length of 4.020*±* 0.126, and an average clustering coefficient of 0.747*±* 006. For each run, two networks were chosen from the 10 networks uniformly at random and one was set as the social network, with the other as the transmission network.

## 3. Results

### 3.1 Model dynamics

We generated time series of the percentage of vaccinators and percentage of infected persons for each of the networks, in order to illustrate the basic dynamics exhibited by the model. We used baseline parameter values everywhere (Table 1) except that *c* = 0.3. For all networks we initialized the population to have a low initial number of vaccinators and a large initial number of susceptible persons. These initial conditions caused the incidence of infection to skyrocket at the beginning of the simulation for all network types (Figure 1). Immediately after this initial outbreak, susceptible neighbours of infected persons get vaccinated, thereby reducing prevalence.

**Figure 1:**
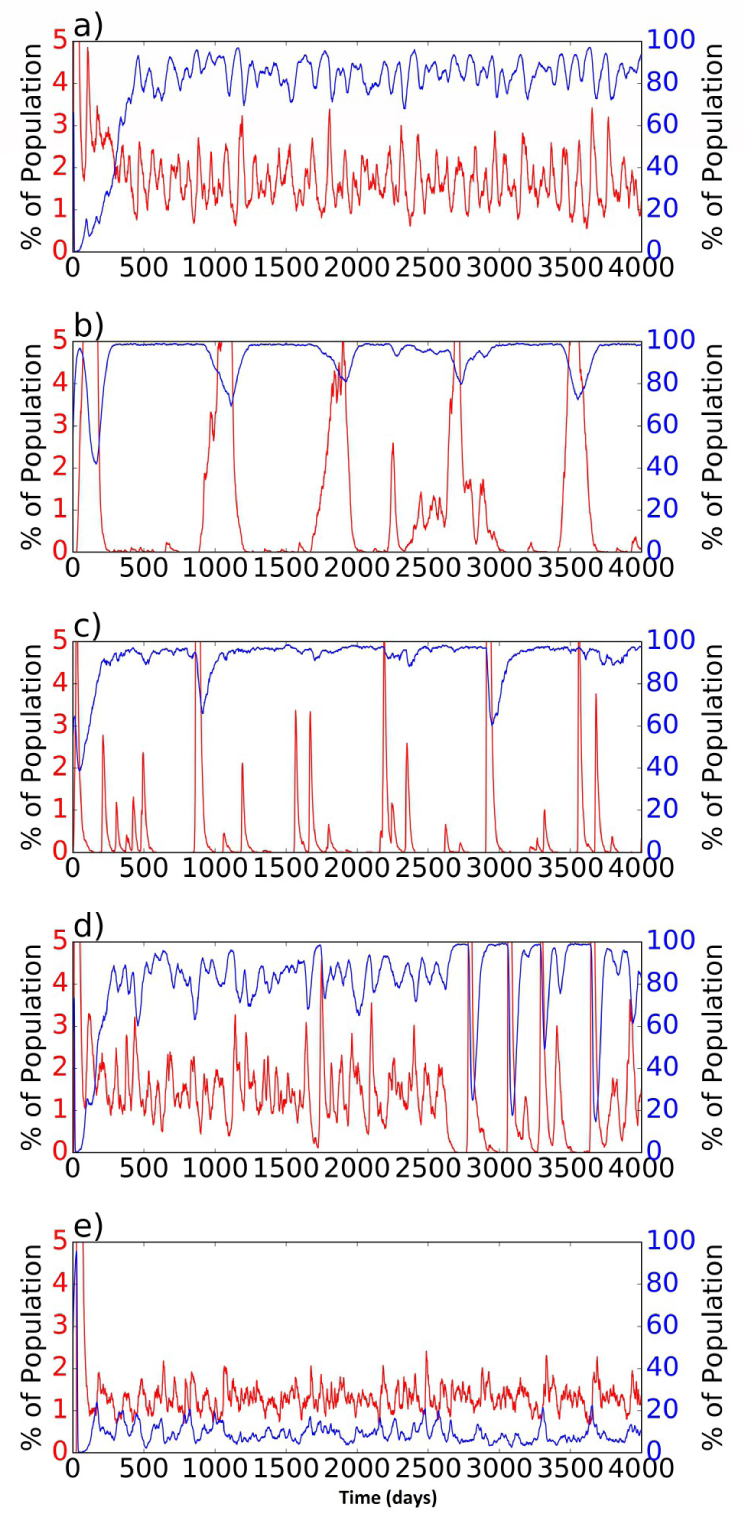
*Time series for a typical simulation on each network type: a) random network, b) square lattice, c) Barabasi-Albert network, d) Small world network, e) empirically-derived networks. Red line is percentage of infected individuals in the population; blue line is percentage of vaccinators in the population. Parameter values are as in Table 1 except c* = 0.3.

After this initial spike, the dynamics settle down into pseudo-stable patterns that vary widely depending on network type. More frequent outbreaks appear to occur on networks with higher degree, which is consistent with intuition (Figure 1). The random network exhibits relatively regular outbreaks (Figure 1a), while the square lattice, Barabasi-Albert network, and small world network exhibit more irregular dynamics consisting of large outbreaks interspersed with periods of very low vaccine coverage and infection prevalence (Figure 1b-d). However, during certain phases in the time series, the small-world network appears to transition to a regime of sustained endemic infection similar to that observed for the random network (Figure 1d). The empirically-derived network exhibits small stochastic fluctuations around an equilibrium, and the percentage of vaccinators is significantly higher in the empirically-derived network than in the other four networks (Figure 1e).

### 3.2 Regime shifts

We carried out this simulation experiment for a range of values of *c* to understand how dynamics respond to changes in the perceived vaccine risk *c*. We computed the long-term average prevalence of infected persons and vaccinators for each value of *c* tested. As *c* approaches zero from below (for *ξ* = 0), a transition from a regime of high vaccine coverage and low infection prevalence to a regime of low vaccine coverage and endemic infection should be observed, since for *c >* 0, the payoff to vaccinate becomes less than the payoff not to vaccinate.

In the simulations we observe a transition in the percentage of nonvaccinators as a function of the perceived vaccine risk *c* in most of the network types (Figure 2). As *c* approaches zero, the prevalence of vaccinators declines dramatically in the first four networks. The transition appears gradual (noncritical) in the empirically-derived network (Figure 2e). We speculate this is due to the greater heterogeneity exhibited by the empirically-derived network than the other four idealized network types. The percentage of infected persons in each network shows similar transitions, even in the latter network (Figure 2e). We also note that the transition is sharper when the sigmoid function used in decision-making is steeper (higher *β*; results not shown).

**Figure 2:**
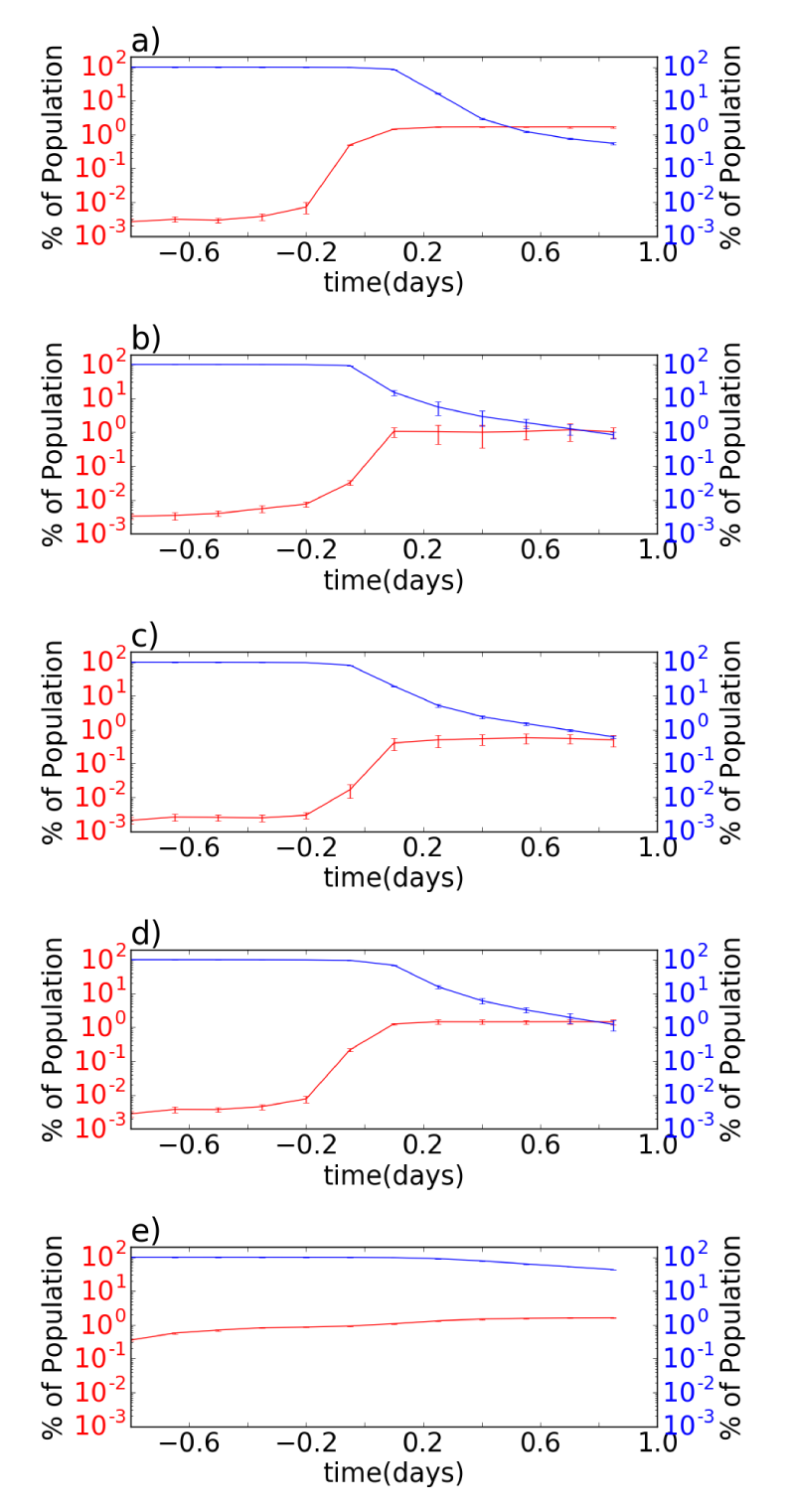
*The time-averaged percentage of infected persons and vaccinators as a function of relative vaccine cost c, showing a critical transition near c* = 0 *on the a) random network,b) square lattice, c) Barabasi-Albert network, d) Small world network, and a more gradual transition on the e) empirically-derived networks. All parameters are as in Table 1 except for c, which is being varied. The blue line represents the percentage of vaccinators, and the red line percentage of infected. Error bars represent the standard deviation over the 100 realizations.*

### 3.3 Early warning signals

Indicators such as spatial correlation can signal an impending critical transition in spatially structured ecological systems [31]. Although theoretical results are not available for coupled behaviour-disease dynamics on multiplex networks, the universality of dynamics near local bifurcations of dynamical systems [32] suggests that similar early warning signals should be observed in our system.

In spatially extended critical phenomena, the plot of spatial correlation versus a bifurcation parameter such as *c* is linear on a log-linear plot [52]. Hence, we computed the average lag-1 spatial correlation (SC) across the entire time series. We repeated this for many values of *c* and plotted lag-1 AC versus *c* on a log-linear scale. As noted previously, we expect near the threshold *c* = 0 where the costs and benefits of the vaccine become balanced, that critical slowing down should emerge in the network, and that this should manifest as increased spatial correlation. As we increase *c* from negative to positive, small clusters of non-vaccinators begin to appear. Each day every node samples a random neighbour, and the only other way for that node to switch opinions is if the randomly sampled neighbour has a different opinion that they do (see Methods). As a result, we expect to see clusters of nonvaccinators emerge, which causes the lag-1 SC to increase before the critical transition (and after which almost everyone because a non-vaccinator) (figure 3).

**Figure 3:**
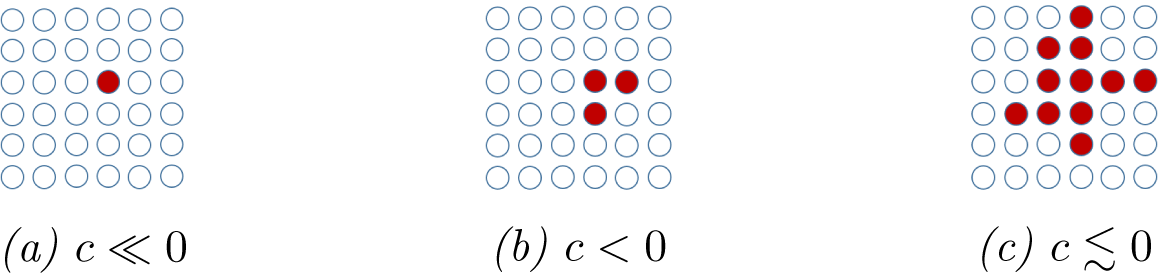
*Visualization of non-vaccinator spatial correlation on a square lattice. As c approaches the critical transition at c* = 0, *clusters of non-vaccinators (red) begin to appear, increasing the spatial correlation of non-vaccinators.*

This pattern is observed in simulations for all network types. As the regime shift at *c* = 0 is approached from negative values of *c* (corresponding to a rise in perceived vaccine risks), we observe a clear and linear increase in the time-averaged lag-1 SC, in plots of the natural logarithm of lag-1 SC versus *c* (Figure 4). This is robust to values of the disease transmission probability, *p* (Figure 4).

**Figure 4:**
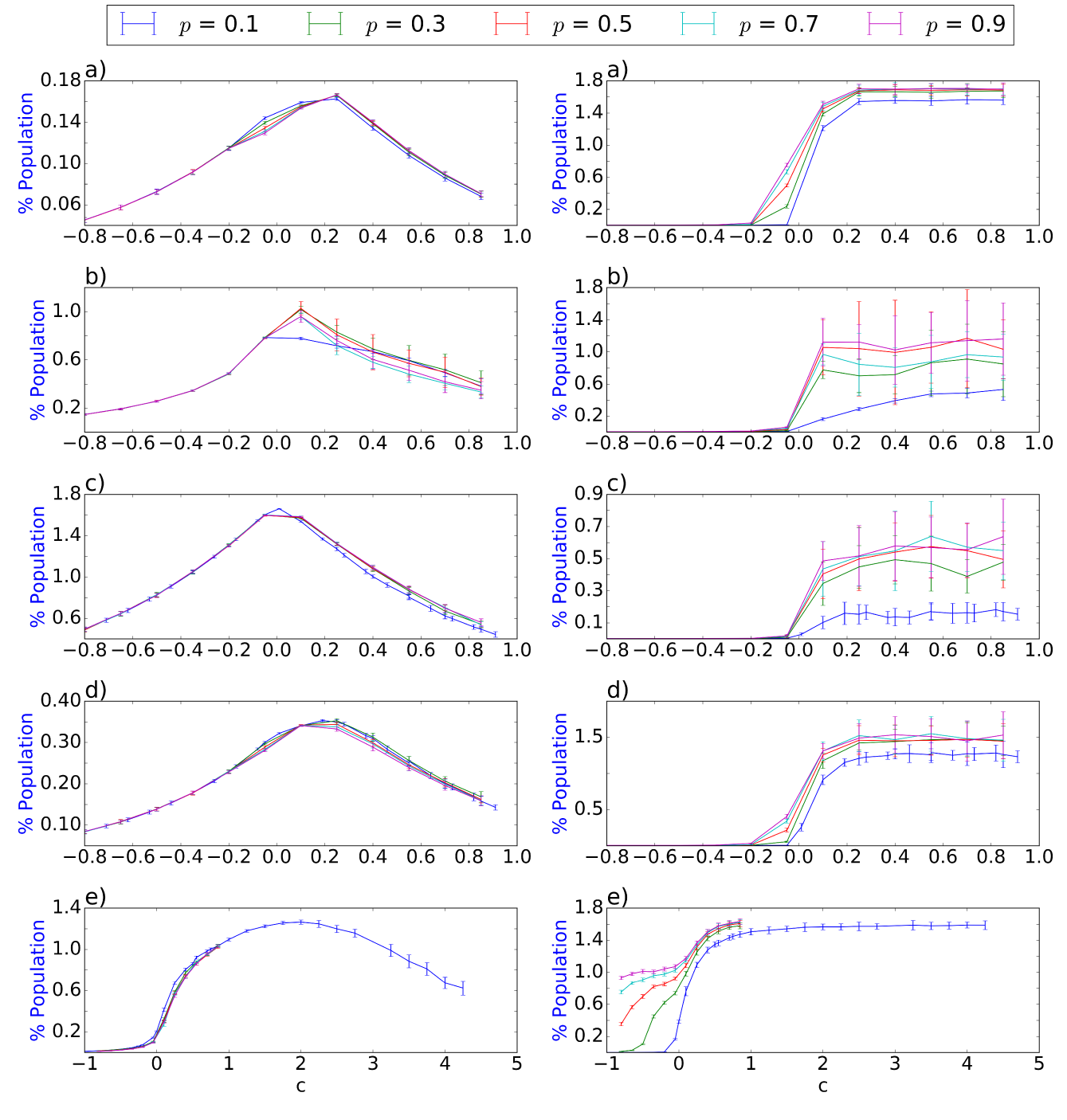
The natural logarithm of the time-averaged lag-1 SC of nonvaccinators, and the percentage of infected nodes, for a range of values of c, showing a linear increase in lag-1 SC in a log-linear plot as the critical transition is approached on a) random network, b) square lattice, c) Barabasi-Albert network, d) Small world network, e) empirically-derived networks. All other parameter values are as in Table 1.

However, there is a notable difference in y-axis scales for the random and small-world networks (Figure 4a,d). Overall these networks show a smaller increase in spatial correlation, possibly due to the smaller average path length in these networks. Furthermore, lag-1 SC in the empirically-derived network has a nonlinear and more gradual response to changes in *c*, which matches the lack of a sharp critical transition in that network. Sensitivity analyses over *r* and *σ* confirm the same patterns, except in the extreme case of r = 0:02 where infected individuals never recover (Figure 5).

**Figure 5:**
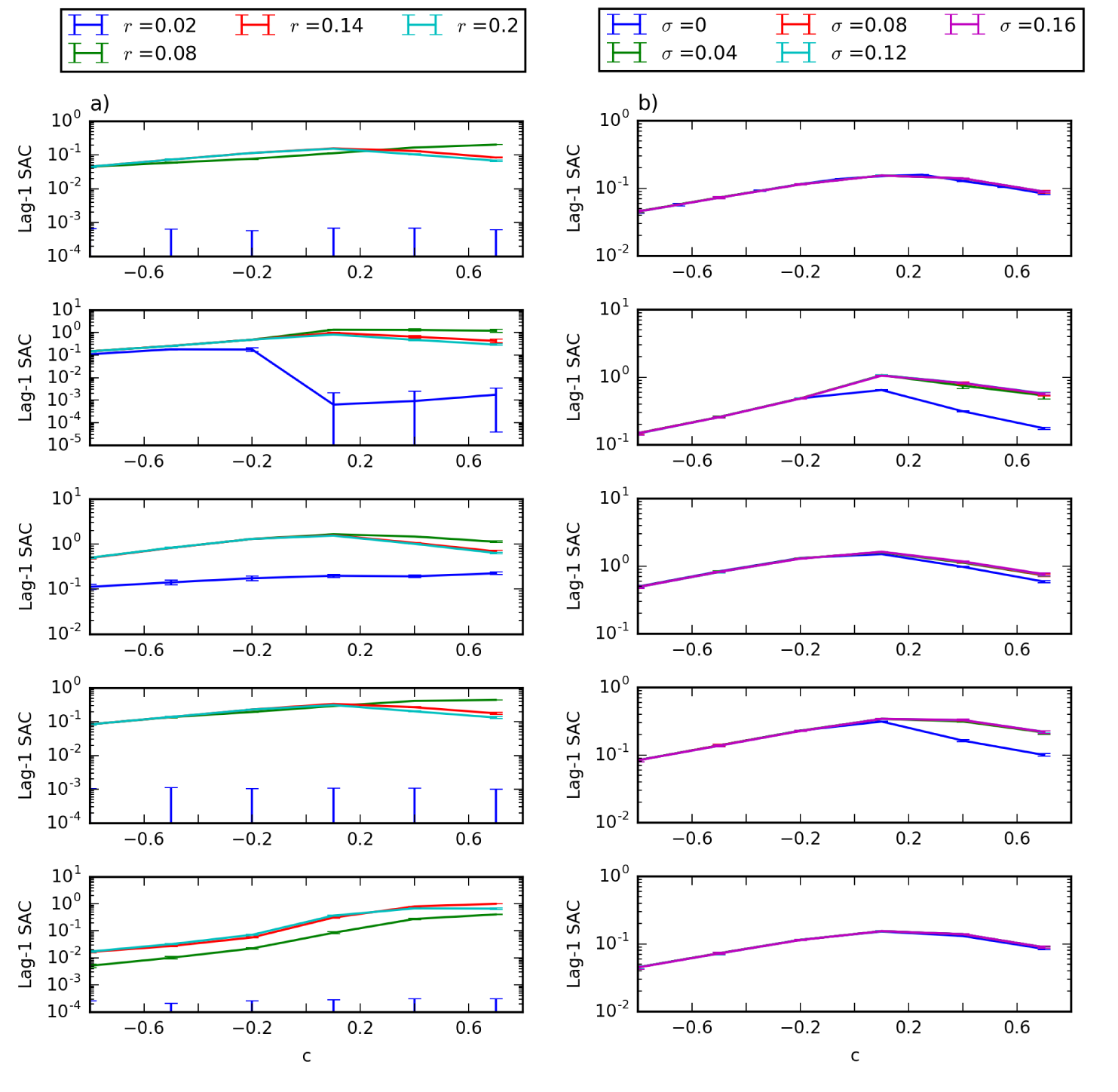
The natural logarithm of the time-averaged lag-1 SC of nonvaccinators for a range of values of c at selected values of a) r and b) σ, showing a linear increase in lag-1 SC in a log-linear plot as the critical transition is approached. Networks types from top row to bottom row are: random network, square lattice, Barabasi-Albert network, small world network, and empirically-derived networks. All parameters besides r, σ and c are the same as Table 1.

We observe that the rise in the natural logarithm of lag-1 SC begins well before the number of non-vaccinators begins to increase appreciably (compare *c ∈* [ 0.8, 0.2] in Figure 4 versus Figure 2). Therefore, tracking lag-1 SC can provide an early warning signals of potential shifts in population vaccinating behaviour that would not be accessible simply by extrapolating the number of non-vaccinators using a linear regression, for instance. Moreover, this rise in lag-1 SC is highly robust to network type and parameter value, due to the fundamental assumption that a node’s vaccination status is influenced by the opinions of the nodes in their social neighbourhood. However, the location of the regime shift in *c* is related to the average node degree: with an average node degree of 100, the regime shift occurs at approximately *c* = 2.4.

## Discussion

Here we studied regime shifts in coupled behaviour-disease dynamics on a multiplex network where an infectious disease is transmitted through the physical network layer, and the social layer describes a population where everyone has either a pro-vaccine or an anti-vaccine opinion. These simulation results show the presence of critical slowing down near a bifurcation in the multiplex network corresponding to a switch from predominant vaccinating behaviour and disease elimination, to predominant non-vaccinating behaviour and disease endemicity. Critical slowing down was clearly manifested in all network types and across a broad range of parameter values, with the exception of the empirically derived network. This exception may have been on account of the greater heterogeneity of the network structure causing lack of a sharp transition to non-vaccinating behaviour.

Hence, the results suggest that it may be possible to use lag-1 spatial correlation in social networks as an early warning signal of widespread vaccine refusal in a population. However, the lack of a clear transition in the case of the network that was empirically derived (from NDSSL data) suggests that further research must be conducted in order to determine how and whether it would be possible to detect such early warning signals in real-world social networks, and what the trends in correlation indicators might signify. We speculate that our approach might have failed for the empirical network due to multiple sources of heterogeneity in network structure such as: a highly dispersed node degree distribution; the presence of disconnected subgraphs; and/or differing network structure in different parts of the network. However, it is possible that including peer pressure (social norms) in the model might cause population opinion states to shift to bistable boundary equilibria corresponding to all-vaccinator or no-vaccinator population compositions–as has been observed in other socio-ecological models–and thus restore the feasibility of early warning signals [26]. Our model also assumed that networks are static and that the two layers are perfectly correlated. Neither condition holds in real populations, and these simplifying assumptions could be relaxed in future work.

It is also possible to tailor this model to specific infectious diseases such as measles or influenza by modifying the model to include relevant vital dynamics, disease natural history, and vaccine characteristics. This is particularly important since disease natural history can have a significant impact on disease dynamics [44, 53], and vaccine coverage can vary widely between both vaccines and populations [54, 55]. Further to this point, there are indications that some disease dynamics, such as meningococcal disease, are in a state of self-evolved criticality in their naturally circulating dynamics (i.e. always close to a critical point) [56]. The impact of ever-present critical disease dynamics on the detectability of early warning signals of a regime shift in a socio-epidemiological state require further research. For instance, the critical disease dynamics could serve to mask early warning signals of socio-epidemiological regime shifts. This would motivate a search for indicators that can distinguish the socio-epidemiological signal from the background of critical disease dynamics.

Finally, future research could seek early warnings signals in lag-1 SC measurements from social networks derived from social media data sources such as Twitter. Lag-1 SC is readily calculated if the sentiment of Twitter users toward vaccines can be assessed as pro- or anti-vaccine. However, the Twitter follower network is a directed graph that changes in time, therefore additional theoretical refinements are necessary. Moreover, our method assumes perfect knowledge of the state of nodes on the social layer, whereas in reality this information is partial. Future work should also explore whether censored data on vaccine opinions changes the reliability of the early warning indicators we explored in this paper. This could be addressed by extended models with a parameter for censoring and a distinction between actual and observed opinion status.

Lag-1 spatial correlation appears to be a robust early warning signal for predicting regime shifts in vaccine uptake under the conditions we studied, indicating potential for worthwhile additional study in the context of coupled behaviour-disease interactions.

## 5 Acknowledgments

The authors are grateful for helpful comments from the editor and reviewers. This research was funded by Natural Sciences and Engineering Research Council of Canada (NSERC) Discovery Grants to MA and CTB.

## References

[1] A. D. Lopez and C. D. Mathers, “Measuring the global burden of disease and epidemiological transitions: 2002–2030,” Annals of tropical medicine and parasitology, 2013.

[2] S. Murch, “Separating inflammation from speculation in autism,” The Lancet, vol. 362, pp. 1498–1499, 2003.

[3] M. Alazraki, “The autism vaccine fraud: Dr. wakefield’s costly lie to society,” Dec 2011.

[4] V. A. Jansen, N. Stollenwerk, H. J. Jensen, M. Ramsay, W. Edmunds, and C. Rhodes, “Measles outbreaks in a population with declining vaccine uptake,” Science, vol. 301, no. 5634, pp. 804–804, 2003.

[5] C. Chen, “Rebellion against the polio vaccine in nigeria: implications for humanitarian policy,” African Health Sciences, vol. 4, pp. 205–207, 2004.

[6] A. S. Jegede, “What led to the Nigerian boycott of the polio vaccination campaign?,” PLoS Med, vol. 4, 2007.

[7] N. H. Fiebach and C. M. Viscoli, “Patient acceptance of influenza vaccination,” The American journal of medicine, vol. 91, no. 4, pp. 393–400, 1991.

[8] W. O. Kermack and A. G. McKendrick, “A contribution to the mathematical theory of epidemics,” Proceedings of the Royal Society of London A: Mathematical, Physical and Engineering Sciences, vol. 115, no. 772, pp. 700–721, 1927.

[9] S. Bansal, B. T. Grenfell, and L. A. Meyers, “When individual behaviour matters: homogeneous and network models in epidemiology,” Journal of the Royal Society Interface, vol. 4, no. 16, pp. 879–891, 2007.

[10] J. Parker and J. M. Epstein, “A distributed platform for global-scale agent-based models of disease transmission,” ACM Transactions on Modeling and Computer Simulation, vol. 22, no. 1, pp. 1–25, 2011.

[11] L. B. Shaw and I. B. Schwartz, “Fluctuating epidemics on adaptive networks,” Physical Review E, vol. 77, no. 6, p. 066101, 2008.

[12] A. Perisic and C. T. Bauch, “Social contact networks and disease eradicability under voluntary vaccination,” PLoS Comput Biol, vol. 5, p. e1000280, 02 2009.

[13] F. Fu, D. I. Rosenbloom, L. Wang, and M. A. Nowak, “Imitation dynamics of vaccination behaviour on social networks,” Proceedings of the Royal Society of London B: Biological Sciences, vol. 278, no. 1702, pp. 42–49, 2011.

[14] S. Funk, E. Gilad, C. Watkins, and V. A. Jansen, “The spread of awareness and its impact on epidemic outbreaks,” Proceedings of the National Academy of Sciences, vol. 106, no. 16, pp. 6872–6877, 2009.

[15] H.-F. Zhang, Z.-X. Wu, M. Tang, and Y.-C. Lai, “Effects of behavioral response and vaccination policy on epidemic spreading-an approach based on evolutionary-game dynamics,” Scientific reports, vol. 4, 2014.

[16] W.-X. Wang, Y.-C. Lai, and C. Grebogi, “Effect of epidemic spreading on species coexistence in spatial rock-paper-scissors games,” Phys. Rev. E, vol. 81, p. 046113, Apr 2010.

[17] A. Perisic and C. T. Bauch, “A simulation analysis to characterize the dynamics of vaccinating behaviour on contact networks,” BMC Infectious Diseases, vol. 9, no. 1, p. 1, 2009.

[18] Z. Wang, M. A. Andrews, Z.-X. Wu, L. Wang, and C. T. Bauch, “Coupled disease-behavior dynamics on complex networks: A review,” Physics of Life Reviews, vol. 15, pp. 1–29, 2015.

[19] E. P. Fenichel, C. Castillo-Chavez, M. G. Ceddia, G. Chowell, P. A. G. Parra, G. J. Hickling, G. Holloway, R. Horan, B. Morin, C. Perrings, and et al., “Adaptive human behavior in epidemiological models,” Proceedings of the National Academy of Sciences, vol. 108, pp. 6306–6311, Apr 2011.

[20] C. T. Bauch and A. P. Galvani, “Social factors in epidemiology,” Science, vol. 342, no. 6154, pp. 47–49, 2013.

[21] C. Granell, S. Gómez, and A. Arenas, “Dynamical interplay between awareness and epidemic spreading in multiplex networks,” Phys. Rev. Lett., vol. 111, p. 128701, Sep 2013.

[22] L. Mao and Y. Yang, “Coupling infectious diseases, human preventive behavior, and networks a conceptual framework for epidemic modeling,” Social Science and Medicine, vol. 74, pp. 167–175.

[23] C. Innes, M. Anand, and C. T. Bauch, “The impact of human-environment interactions on the stability of forest-grassland mosaic ecosystems,” Scientific reports, vol. 3, p. 2689, 2013.

[24] L.-A. Barlow, J. Cecile, C. T. Bauch, and M. Anand, “Modelling interactions between forest pest invasions and human decisions regarding firewood transport restrictions,” PLoS One, vol. 9, no. 4, p. e90511, 2014.

[25] K. A. Henderson, C. T. Bauch, and M. Anand, “Alternative stable states and the sustainability of forests, grasslands, and agriculture,” Proceedings of the National Academy of Sciences, vol. 113, no. 51, pp. 14552–14559, 2016.

[26] R. P. Sigdel, M. Anand, and C. T. Bauch, “Competition between injunctive social norms and conservation priorities gives rise to complex dynamics in a model of forest growth and opinion dynamics,” Journal of theoretical biology, vol. 432, pp. 132–140, 2017.

[27] Q. Guo, X. Jiang, Y. Lei, M. Li, Y. Ma, and Z. Zheng, “Two-stage effects of awareness cascade on epidemic spreading in multiplex networks,” Phys. Rev. E, vol. 91, p. 012822, Jan 2015.

[28] S. Xia and J. Liu, “A computational approach to characterizing the impact of social influence on individuals vaccination decision making,” PLoS ONE, vol. 8, p. e60373, 04 2013.

[29] M. Scheffer, J. Bascompte, W. A. Brock, V. Brovkin, S. R. Carpenter, V. Dakos, H. Held, E. H. V. Nes, M. Rietkerk, and G. Sugihara, “Early-warning signals for critical transitions,” Nature, vol. 461, pp. 53–59, 2009.

[30] C. T. Bauch, R. Sigdel, J. Pharaon, and M. Anand, “Early warning signals of regime shifts in coupled human–environment systems,” Proceedings of the National Academy of Sciences, p. 201604978, 2016.

[31] V. Dakos, E. van Nes, R. Donangelo, H. Fort, and M. Scheffer, “Spatial correlation as leading indicator of catastrophic shifts,” Theoretical Ecology, vol. 3, no. 3, pp. 163–174, 2010.

[32] C. Boettiger, N. Ross, and A. Hastings, “Early warning signals: the charted and uncharted territories,” Theoretical ecology, vol. 6, no. 3, pp. 255–264, 2013.

[33] M. Scheffer, V. Dakos, and E. H. V. Nes, “Slowing down as an early warning signal for abrupt climate change,” IOP Conference Series: Earth and Environmental Science, vol. 105, pp. 14308 – 14312, 2009.

[34] C. E. Elger and K. Lehnertz, “Seizure prediction by non-linear time series analysis of brain electrical activity,” European Journal of Neuroscience, vol. 10, pp. 786–789, 1998.

[35] B. Lebaron, “Some relations between volatility and serial correlations in stock market returns,” The Journal of Business, vol. 65, pp. 199–199, 1992.

[36] S. R. Carpenter and W. A. Brock, “Early warnings of regime shifts in spatial dynamics using the discrete fourier transform,” Ecosphere, vol. 1, pp. 2150–8925, 2010.

[37] T. J. Cline, D. A. Seekell, S. R. Carpenter, M. L. Pace, J. R. Hodgson, J. F. Kitchell, and B. C. Weidel, “Early warnings of regime shifts: evaluation of spatial indicators from a whole-ecosystem experiment,” Ecosphere, vol. 5, no. 8, 2014.

[38] V. Guttal and C. Jayaprakash, “Spatial variance and spatial skewness: leading indicators of regime shifts in spatial ecological systems,” Theoretical Ecology, vol. 2, no. 1, pp. 3–12, 2009.

[39] C. R. Wells, E. Y. Klein, and C. T. Bauch, “Policy resistance undermines superspreader vaccination strategies for influenza,” PLoS Comput Biol, vol. 9, p. e1002945, 03 2013.

[40] A. Okabe and K. Sugihara, Spatial analysis along networks: statistical and computational methods. Wiley, 2012.

[41] “Spatial correlation at lag 1,” Early Warning Signals Toolbox, 2015.

[42] S. Kefi, V. Guttal, W. A. Brock, S. R. Carpenter, A. M. Ellison, V. N. Livina, D. A. Seekell, M. Scheffer, E. H. van Nes, and V. Dakos, “Early warning signals of ecological transitions: Methods for spatial patterns,” PLoS ONE, vol. 9, p. e92097, 03 2014.

[43] V. Dakos, S. Kefi, M. Rietkerk, E. H. V. Nes, and M. Scheffer, “Slowing down in spatially patterned ecosystems at the brink of collapse,” The American Naturalist, vol. 177, no. 6, 2011.

[44] C. T. Bauch and D. J. Earn, “Transients and attractors in epidemics,” Proceedings of the Royal Society of London B: Biological Sciences, vol. 270, no. 1524, pp. 1573–1578, 2003.

[45] S. Bansal, B. Pourbohloul, and L. A. Meyers, “A comparative analysis of influenza vaccination programs,” PLoS Med, vol. 3, no. 10, p. e387, 2006.

[46] A. E. Fiore, D. K. Shay, K. Broder, J. K. Iskander, T. M. Uyeki, G. Mootrey, J. S. Bresee, and N. J. Cox, “Centers for disease control and prevention,” Aug 2008.

[47] D. M. Vickers, A. M. Anonychuk, P. De Wals, N. Demarteau, and C. T. Bauch, “Evaluation of serogroup c and acwy meningococcal vaccine programs: Projected impact on disease burden according to a stochastic two-strain dynamic model,” Vaccine, vol. 33, no. 1, pp. 268–275, 2015.

[48] P. Erdos and A. Renyi, “On random graphs,” Publicationes Mathematicae, vol. 6, pp. 290–297, 1959.

[49] R. Albert and A.-L. Barabasi, “On random graphs,” Science, vol. 286, pp. 509–512, 1999.

[50] D. Easley and J. Kleinberg, Networks, crowds, and markets reasoning about a highly connected world. Cambridge University Press, 2010.

[51] “Synthetic data products for societal infrastructures and proto-populations: Data set 1.0,”

[52] D. Ivaneyko, J. Ilnytskyi, B. Berche, and Y. Holovatch, “Local and cluster critical dynamics of the 3d random-site ising model,” Physica A: Statistical Mechanics and its Applications, vol. 370, no. 2, pp. 163–178, 2006.

[53] J. Dushoff, J. B. Plotkin, S. A. Levin, and D. J. Earn, “Dynamical resonance can account for seasonality of influenza epidemics,” Proceedings of the National Academy of Sciences of the United States of America, vol. 101, no. 48, pp. 16915–16916, 2004.

[54] T. A. e. a. Santibanez, “Flu vaccination coverage, united states, 2014-15 influenza season,” 2015.

[55] L. D. Elam-Evans, D. Yankey, J. Singleton, and M. Kolasa, “National, state, and selected local area vaccination coverage among children aged 19-35 months, United States, 2013,” 2014.

[56] N. Stollenwerk, M. C. Maiden, and V. A. Jansen, “Diversity in pathogenicity can cause outbreaks of meningococcal disease,” Proceedings of the National Academy of Sciences of the United States of America, vol. 101, no. 27, pp. 10229–10234, 2004.

